# A cloud-based workflow to quantify transcript-expression levels in public cancer compendia

**DOI:** 10.1101/063552

**Authors:** PJ Tatlow, Stephen R. Piccolo

## Abstract

Public compendia of raw sequencing data are now measured in petabytes. Accordingly, it is becoming infeasible for individual researchers to transfer these data to local computers. Recently, the National Cancer Institute funded an initiative to explore opportunities and challenges of working with molecular data in cloud-computing environments. With data in the cloud, it becomes possible for scientists to take their tools to the data and thereby avoid large data transfers. It also becomes feasible to scale computing resources to the needs of a given analysis. To evaluate this concept, we quantified transcript-expression levels for 12,307 RNA-Sequencing samples from the Cancer Cell Line Encyclopedia and The Cancer Genome Atlas. We used two cloud-based configurations to process the data and examined the performance and cost profiles of each configuration. Using “preemptible virtual machines”, we processed the samples for as little as $0.09 (USD) per sample. In total, we processed the TCGA samples (n=11,373) for only $1,065.49 and simultaneously processed thousands of samples at a time. As the samples were being processed, we collected detailed performance metrics, which helped us to track the duration of each processing step and to identify computational resources used at different stages of sample processing. Although the computational demands of reference alignment and expression quantification have decreased considerably, there remains a critical need for researchers to optimize preprocessing steps (e.g., sorting, converting, and trimming sequencing reads). We have created open-source *Docker* containers that include all the software and scripts necessary to process such data in the cloud and to collect performance metrics. The processed data are available in tabular format and in Google's BigQuery database (see https://osf.io/gqrz9).

## Introduction

Over the past decade, public cancer compendia have played a crucial role in enabling scientists to identify genomic, transcriptomic, proteomic, and epigenomic factors that influence tumor initiation, progression, and treatment responses^1–19^. Due to efforts like The Cancer Genome Atlas (TCGA), International Cancer Genomics Consortium, Cancer Cell Line Encyclopedia (CCLE), and Connectivity Map, thousands of studies have been published. Typically, consortia who oversee these efforts release raw *and* preprocessed data for the public to use. Accordingly, researchers who wish to reprocess raw data using alternative methods may do so^20–22^. For example, we previously reprocessed 10,005 RNA-Sequencing samples from TCGA and demonstrated that an alternative pipeline provided analytical advantages over the preprocessed data provided by the TCGA consortium^20^. However, this effort required us to copy more than 50 terabytes of data, across three time zones, from the data repository to our local file servers—and to employ tens of thousands of hours of computational time on local computer clusters. Other efforts, such as the Genomic Data Commons^23^, are also reprocessing cancer compendia using updated pipelines. Such efforts require considerable institutional investment in computational infrastructure^24^. In the case of raw sequencing data, computational infrastructure must also implement appropriate security measures to ensure patient privacy^25,26^. Many research institutions do not have the resources to support such infrastructure, and duplicate efforts may unwittingly occur, resulting in wasted resources.

As an alternative, the National Cancer Institute initiated the Cancer Genomics Cloud Pilots^27^, which enable researchers to access cancer compendia via cloud-computing services, such as Google Cloud Platform^28^ or Amazon Web Services^29^. Via these public/private partnerships, cancer data are stored (and secured, as necessary) in shared computing environments. Researchers can rent virtual machines (VMs) in these environments and apply computational tools to the data, without needing to transfer the data to or from an outside location. This model promises to speed the process of scientific discovery, reduce barriers to entry, and democratize access to the data^30,31^. In the current era of highly collaborative science, this model also makes it easier for researchers from multiple institutions to collaborate in the same computing environment.

The landscape of bioinformatics tools available to process RNA-Sequencing data is rapidly evolving. Seemingly small differences in software versions or annotations can lead to considerable analytical differences or make it difficult to integrate datasets^32,33^. However, because the Cancer Genomics Cloud Pilots provide access to raw data, researchers can reprocess the data using whatever tools and annotations support their specific needs.

In coordination with the Institute for Systems Biology (ISB)^34^, we used the Google Cloud Platform to process 12,307 RNA-Sequencing samples from the CCLE and TCGA projects. After preprocessing, we aligned the sequencing reads to the most current GENCODE reference transcriptome (see Methods) and calculated transcript-expression levels using *kallisto*, a pseudoalignment and read-quantification program that executes considerably faster than previous-generation tools, while attaining similar levels of accuracy^35^. We encapsulated all the software necessary to perform these steps into software containers^33,36^. Where possible, the containers execute tasks in parallel and dynamically determine the number of tasks that can be executed simultaneously. In addition, the containers collect detailed computer-performance metrics while the containers execute. We have made these containers freely available to the research community—along with scripts for executing and monitoring them (see Methods).

We processed the CCLE samples (n = 934) using either 1) a cluster-based configuration, orchestrated by the *Kubernetes* system^37^, or 2) preemptible VMs. In this paper, we describe our experiences with these deployment approaches. The cluster-based configuration more closely resembles computing environments typically available at research institutions and thus may be more intuitive for researchers to use. However, using preemptible VMs, we were able to process the data at a considerably lower cost and with less monitoring overhead. Therefore, we used preemptible VMs to process all available TCGA RNA-Sequencing samples (n=11,373) for a total cost of $1,065.49.

Below we describe lessons learned as we processed these data, and we discuss logistical and financial issues that should be considered when using cloud-computing environments. We hope these observations will enable researchers to better evaluate options for processing large biological datasets in the cloud. We also explore opportunities for bioinformaticians to optimize data processing.

## Results

### Cluster-based configuration (CCLE data)

We created a software container to quantify transcript-expression levels for 934 RNA-Sequencing samples from CCLE on the Google Cloud Platform. Initially, we processed these samples on a cluster of 295 computing nodes. Each computing node had access to 4 virtual central processing units (vCPUs), 26 gigabytes of random access memory (RAM), and 400 gigabytes of disk-storage space. We used the *Kubernetes* system to distribute execution across these nodes in a queue-like manner: as one sample completed processing, a new sample began processing, and so on.

The CCLE samples were available in BAM format^38^, whereas *kallisto* only accepts input files in FASTQ format. Therefore, the first step in our pipeline was to sort the BAM data by name and convert the data to FASTQ format (see Methods). To enable faster execution, we performed these steps in parallel (see Methods). On average, these preprocessing steps took 84.7 minutes per sample (71.7% of the processing time), whereas *kallisto* took only 29.8 minutes (25.2% of the processing time; see Figure 1). The remaining 3.6 minutes (3.0%) were used to copy files to and from the computing nodes. These observations indicate that RNA-Sequencing preprocessing steps require a considerable amount of computational time and must be taken into account when planning resource allocation and estimating costs.

One limitation of a cluster-based configuration is that all computing nodes must remain active while even one sample is still processing. In general, larger RNA-Sequencing samples take longer to process than smaller samples. Therefore, in an attempt to reduce costs, we submitted samples to the queue in order of size (largest to smallest). Figure 2 illustrates the start and end times at which each sample was processed. The longest-running sample finished processing in 274 minutes, while one sample took only 46 minutes. Few computing nodes were idle during the first 350 minutes of overall processing time, and all but two samples had completed processing after 450 minutes. Disk-storage drives failed to mount properly while the remaining two samples were processing, so these samples had to be resubmitted to the queue.

**Figure 1.**
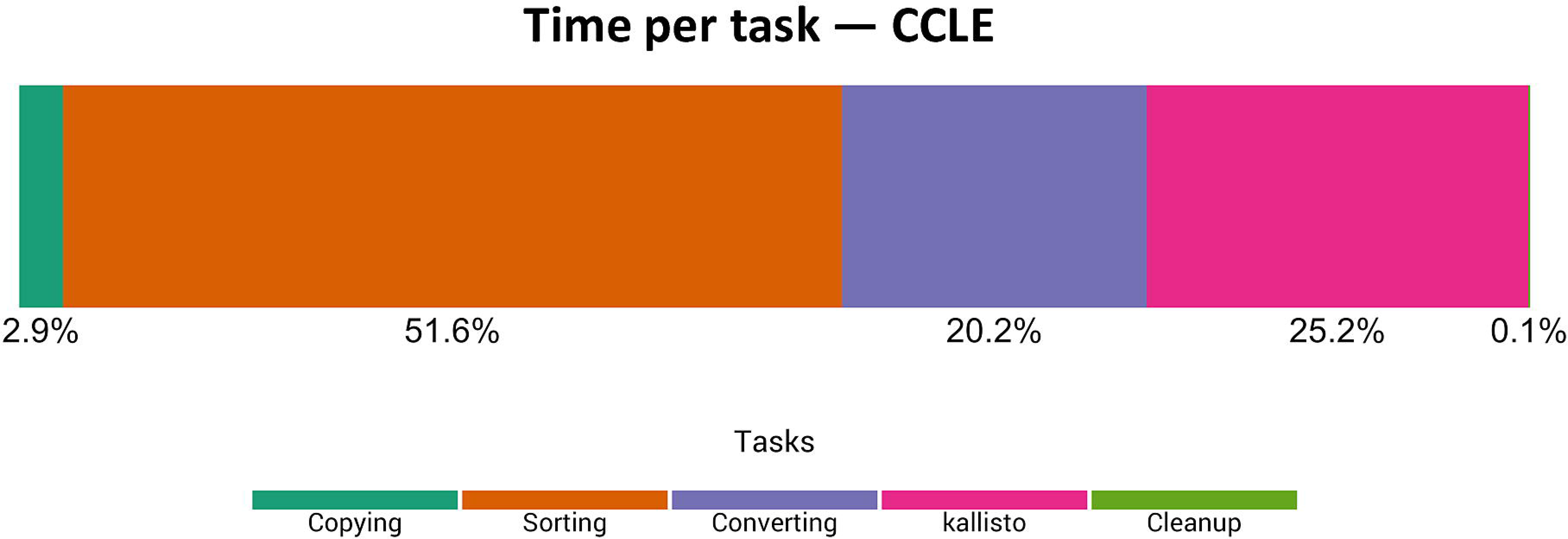
Relative time spent on computational tasks for CCLE samples using the cluster-based configuration. We logged the durations of individual processing tasks for all CCLE samples, averaged these values, and calculated the percentage of overall processing time for each task. Because the raw data were stored on Google Cloud Storage, copying the BAM and index files to the computing nodes took less than 3% of the total processing time. For preprocessing, the BAM files were sorted and converted to FASTQ format, which took 71.8% of the overall processing time. The *kallisto* alignment and quantification steps took only 25.2% of the overall processing time.

**Figure 2.**
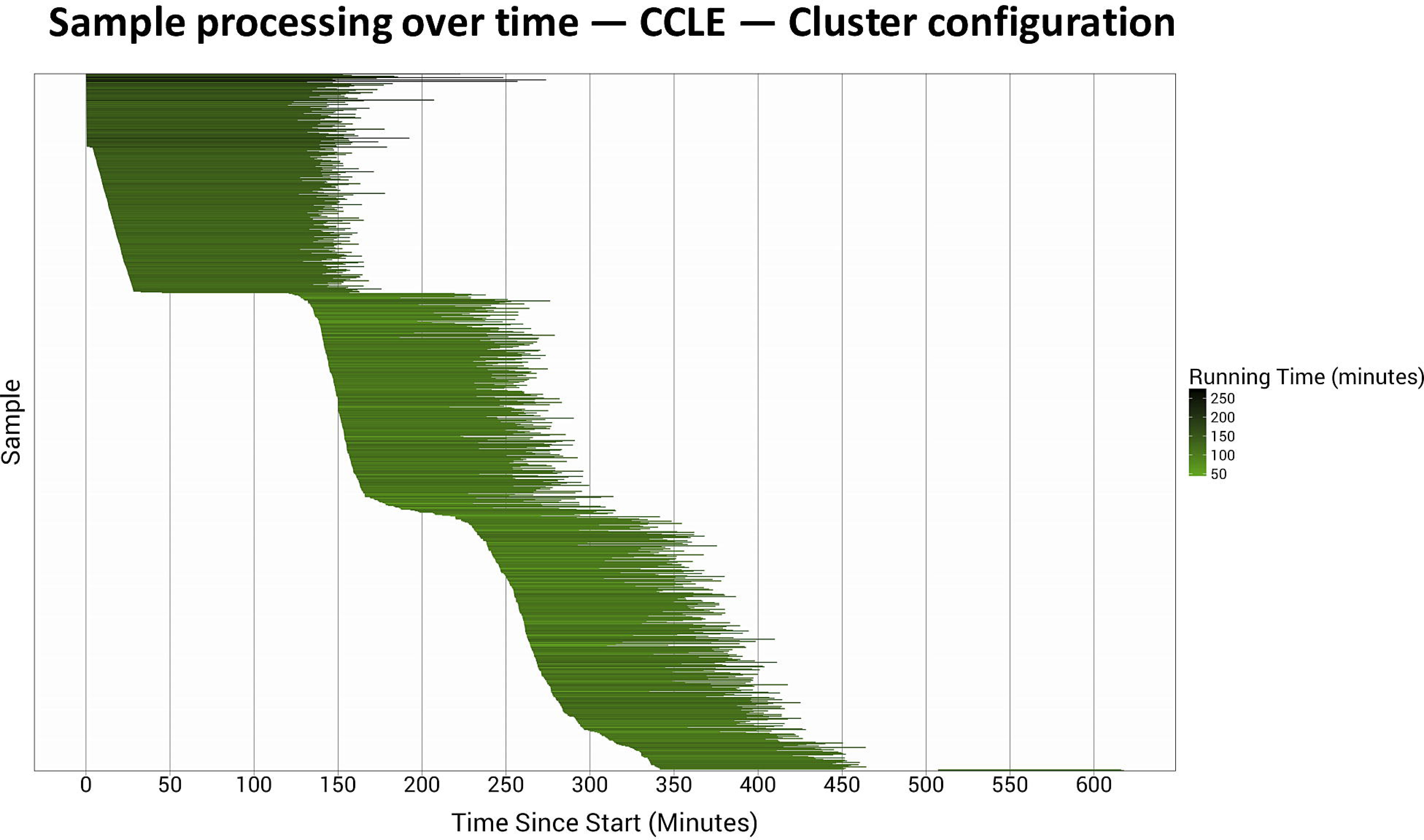
Processing time per CCLE sample using the cluster-based configuration. The 934 CCLE samples were processed on a cluster of 295 virtual machines. The horizontal lines represent the relative start and stop times at which each sample was processed. Darker lines identify samples that took longer to process. The longest-running sample took over 4.5 hours to process; the shortest-running sample completed in less than 1 hour. All but two samples finished processing within 7.5 hours. These two samples failed due to a disk-mounting error, so we reprocessed the samples on a smaller cluster (see Methods).

We collected computer-performance metrics at one-second intervals while the software container was executing. Resource utilization varied dramatically across the processing tasks (Figure 3). The sorting and *kallisto* phases were relatively vCPU intensive, whereas the BAM-to-FASTQ conversion used the disk-storage devices relatively heavily. RAM usage was consistently high, although it decreased temporarily during the sorting and conversion stages. As expected, network usage was high while data files were being transferred to and from computing nodes—but zero at other times. vCPU usage was highly variable across the individual vCPUs on a given computing node, and vCPUs were never used at full capacity for a considerable length of time.

**Figure 3.**
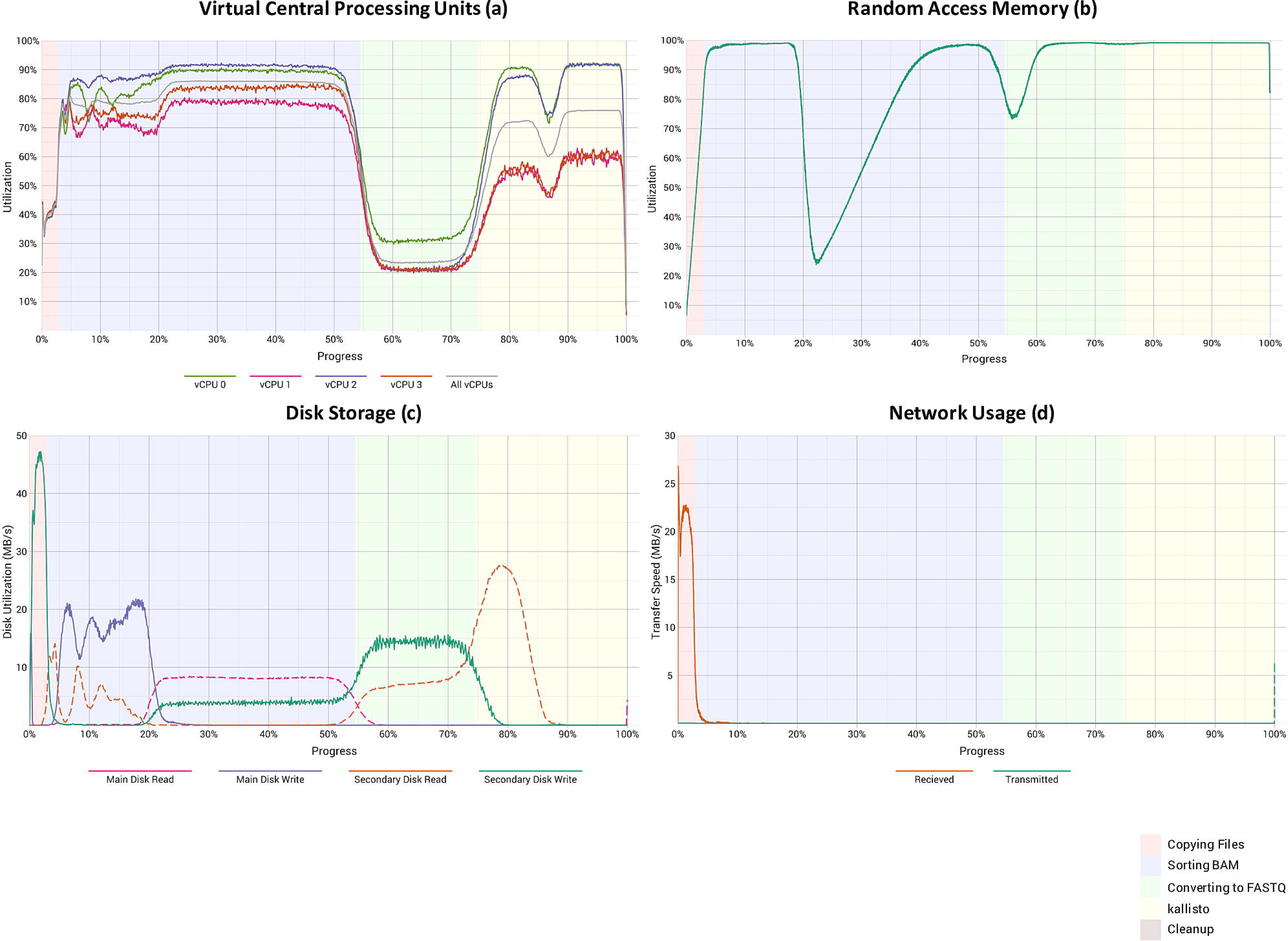
Computational resource utilization while CCLE samples were processed using a cluster-based configuration. These graphs show the (a) percentage of user and system vCPU utilization, (b) percentage of memory usage, (c) disk activity, and d) network activity. The “main” disk for each virtual machine had 100 gigabytes of storage space. The “secondary” disks, which stored temporary files during the sorting and FASTQ-to-BAM conversion steps, had 300 gigabytes of space. The background colors represent the five computational tasks shown in Figure 2. Each graph summarizes data from all 934 CCLE samples.

The cost for each node in the cluster was $0.252 (USD) per hour. There was no cost for the master node that orchestrated the cluster. These computing costs and data-storage costs constituted the great majority of expenses that we incurred (see Tables 1-2). In total, the cost was $716.52 ($0.77 per sample). Because cloud-computing offerings, service prices, and bioinformatics tools change regularly, it is difficult to compare costs across projects. However, a 2011 study used the MAQ software^39^ to align a single biological sample in the cloud for $320.10^29^.

**Table 1:** Summary of data sets and computational resources.

| Category | CCLE<br>(cluster) | CCLE<br>(preemptible) | TCGA<br>(breast, lung) | TCGA (other<br>cancer types) |
| --- | --- | --- | --- | --- |
| # samples | 934 | 934 | 1,811 | 9,562 |
| Data input format | BAM | BAM | FASTQ | FASTQ |
| Total data size<sup>a</sup> | 12.59 | 12.59 | 10.67 | 57.8 |
| # vCPUs per node<sup>b</sup> | 4 | 16 | 2 | 2 |
| RAM per node<sup>c</sup> | 26 | 104 | 7.5 | 7.5 |
| Primary disk storage per<br>node<sup>d</sup> | 100 | 100 | 10 | 10 |
| Secondary disk storage<br>per node<sup>e</sup> | 300 | 300 | 350 | 350 |

<sup>a</sup> Total amount of raw sequencing data (measured in terabytes).
<sup>b</sup> vCPU = virtual central processing unit.
<sup>c</sup> RAM = random access memory (measured in gigabytes).
<sup>d</sup> The primary disks were used to store operating system components and temporary files.
Storage capacity was measured in gigabytes.
<sup>e</sup> The secondary disks were used to store input and output data.

### Preemptible-node configuration (CCLE data)

Next we reprocessed the CCLE RNA-Sequencing samples using an alternative approach. First, to assess whether quality trimming would have a significant effect on the *kallisto* quantification values, we excluded low-quality bases and any adapter sequences that were present in the data. Second, we used preemptible VMs and compared the performance and cost profiles of this configuration against what we observed for the cluster-based option.

Typically, cloud-service providers have more computing nodes available than customers will use at any given time. While these extra nodes are idle, users can request to use the nodes with “preemptible” status—at a considerably reduced price. When demand rises, the service provider halts (preempts) execution of the nodes and makes them available to full-paying customers. In situations where nodes are preempted infrequently—and preemptible nodes are available—this approach can reduce costs, although additional care must be taken to ensure that preempted samples are re-executed when new nodes become available.

To reduce the time that the CCLE samples would take to process—and thus to reduce the likelihood of preemption—we used more powerful computing nodes: 16 vCPUs and 104 gigabytes of RAM per node. As the samples were processing, we executed a script, every 60 seconds, to check whether any samples had been preempted and then resubmitted those samples for processing (see Methods). Unlike the cluster-based configuration, which employed a static number of computing nodes, the number of preemptible nodes changed dynamically, according to availability. During the first 20 minutes, 932 nodes were allocated, and most samples completed processing before 150 total minutes had elapsed (Figure S1). Of the 934 CCLE samples, 78 were preempted at least once (Figure S1). Eight samples were preempted twice, and one sample was preempted three times. On average, preempted nodes executed for 76.64 minutes (min = 1.87; max = 126.57) before preemption. The BAM files for the preempted samples averaged 14.07 gigabytes in size, whereas the remaining samples averaged 13.79 gigabytes. Accordingly, preemption was a relatively random process, not necessarily influenced by the size of the data files.

Trimming next-generation sequencing reads is a common, though somewhat controversial, practice^40–42^. Trimming can remove low-quality bases and reads and can remove artefactual adapter sequences. We compared transcript-expression levels from our initial (untrimmed) CCLE data against the data that had been subjected to trimming. The average, sample-wise Spearman correlation coefficient was 0.988; however, the correlation coefficients for some samples were as low as 0.92 (Figure S2), suggesting that these samples had a considerable number of low-quality reads.

The performance profiles for the preemptible configuration were similar to those from the cluster-based configuration (Figures S3-S4). The read-trimming step required an average of 7.97 minutes (9.0%) of processing time per sample (see Figure S3). In addition, there was considerable overhead to “spin up” and configure the VMs and to localize the necessary files (19.4% of processing time). We suspect this overhead was relatively high because the cloud environment needed to allocate a relatively large amount of resources (vCPUs and RAM) per computing node. Overall, the *kallisto* alignment and quantification process took only 9.8% of the processing time, reaffirming that preprocessing steps may contribute more to the expense of processing RNA-Sequencing data than alignment and quantification, which have been optimized substantially in recent years.

The cost of the preemptible nodes (16 vCPU) was $0.280 per hour. In addition, we rented a master node for $0.01 per hour. The total cost to process the samples was $364.22 ($0.39 per sample). Even though preemption occurred 88 times—contributing an estimated $31.48 to the total cost—the overall cost was 49.2% lower than the cluster-based configuration. This was true, even though we added the read-trimming step. Accordingly, we conclude that preemptible nodes show promise as a way to reduce expenses markedly, so long as preemption rates remain low.

### Preemptible-node configuration (TCGA data)

Based on our experiences with the CCLE samples, we used preemptible VMs to process 11,373 RNA-Sequencing samples from TCGA that spanned 34 cancer types. The TCGA samples were available in FASTQ format; therefore, we did not need to sort or convert the data. However, the FASTQ file were packed inside “tar” files, so we included a step to unpack these files. Based on preliminary testing, we estimated that using less-powerful computing nodes—2 vCPUs and 7.5 gigabytes of RAM—would provide an optimal balance between cost and performance.

We started with 1,811 breast carcinoma and lung squamous cell carcinoma samples. These samples completed processing in 101 minutes on average, compared to 88.5 minutes for the CCLE samples. Although individual TCGA samples took 14% longer to process than the CCLE samples, the computing nodes used 87.5% fewer computational resources per sample. The VM “spinup” and configuration tasks were considerably shorter (Figure 4), ostensibly because the computing nodes required fewer resources. The *kallisto* step took 41.3% of the processing time, due to the shorter preprocessing times. Only 210 preemptions occured (Figure 5).

**Figure 4.**
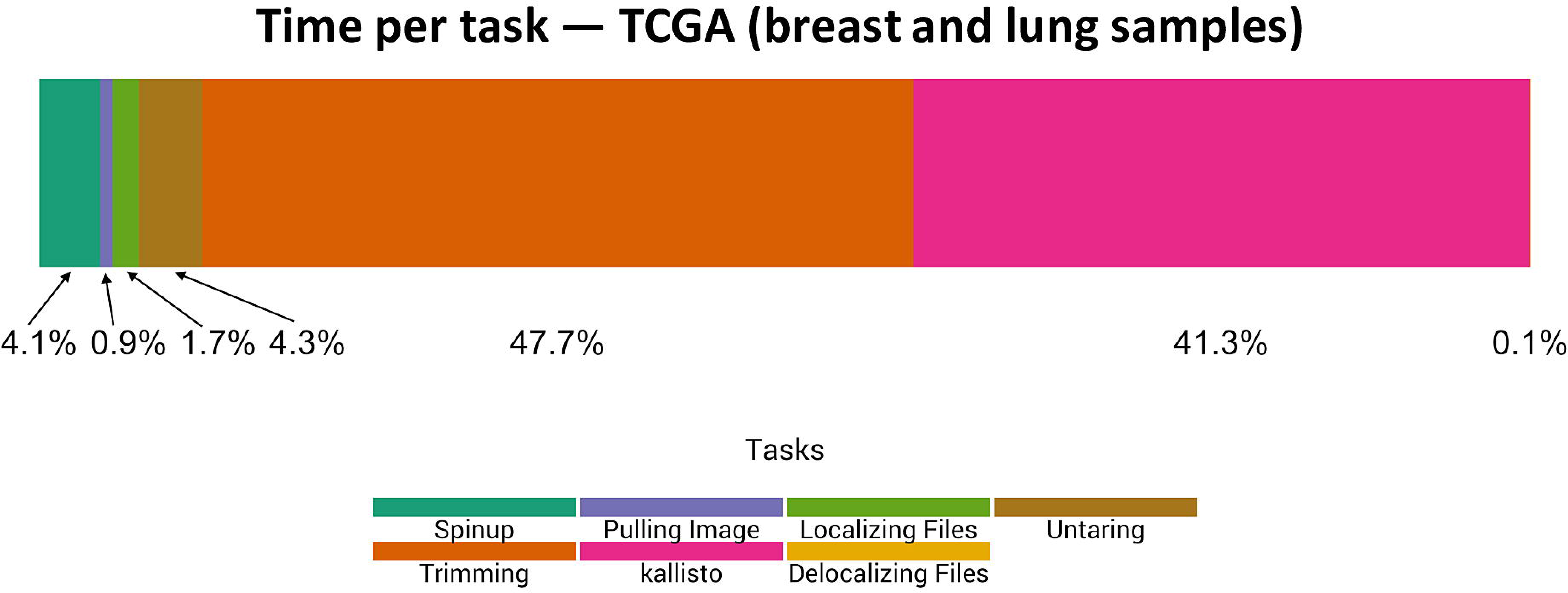
Relative time spent on computational tasks for TCGA breast and lung samples using the preemptible-node configuration. We logged the durations of individual processing tasks for the TCGA breast and lung samples, averaged these values, and calculated the percentage of overall processing time for each task. The “spinup,” image pulling, and file localization steps enabled the virtual machines to begin executing. For sample preprocessing, the FASTQ files were unpacked, decompressed, and quality trimmed; together these steps took 52.0% of the processing time (on average). The *kallisto* alignment and quantification steps took 41.3% of the overall processing time.

**Figure 5.**
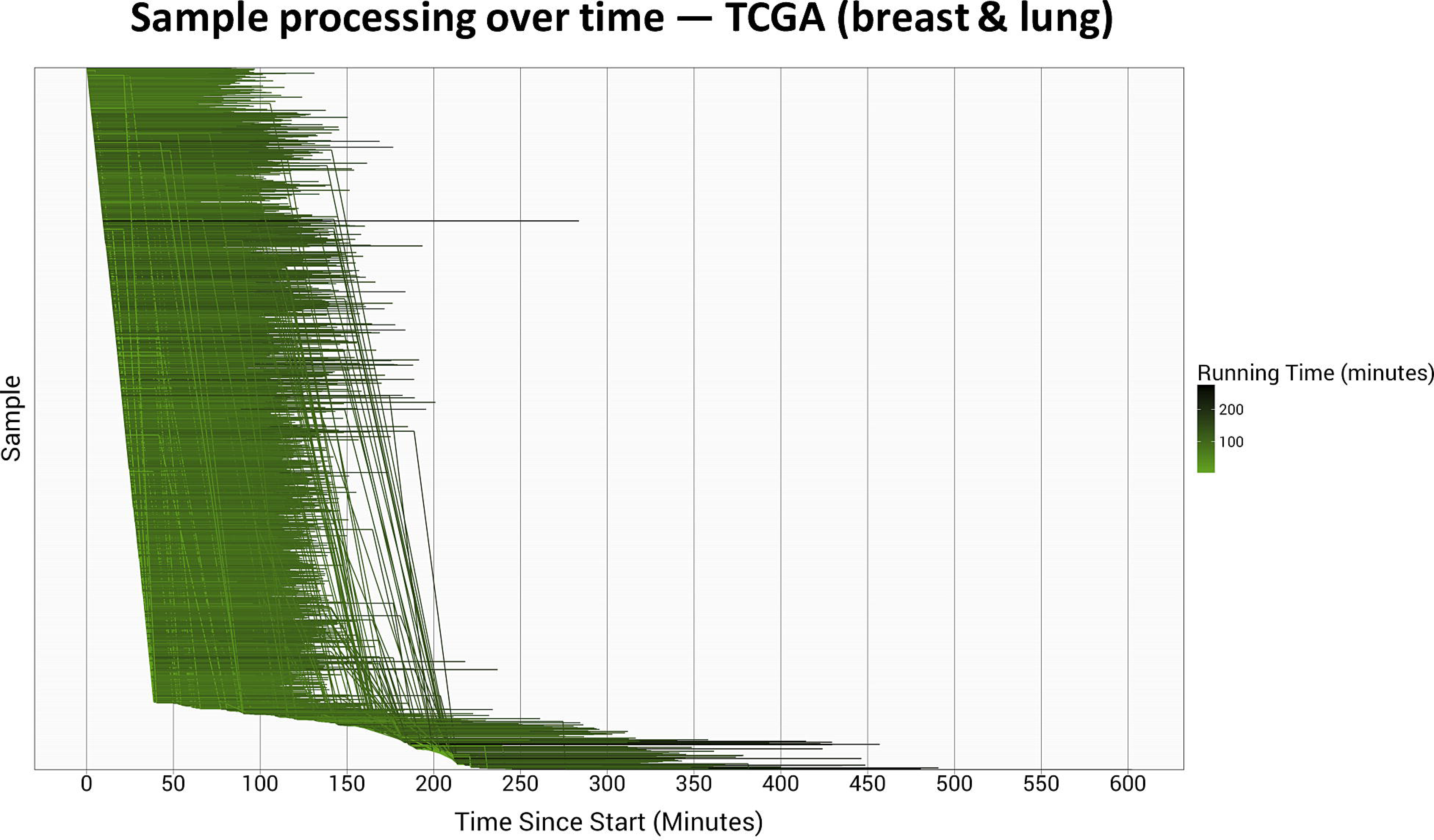
Processing time per TCGA breast and lung sample using the preemptible-node configuration. The 1,811 TCGA breast and lung samples were processed using a variable number of preemptible virtual machines. The horizontal lines represent the relative start and stop times at which each sample was processed. Darker lines identify samples that took longer to process. Vertical lines indicate times at which samples were preempted and then resubmitted for processing. In total, 210 preemptions occurred.

Because most of the TCGA samples were assigned to a single read group, we were not able to parallelize the quality-trimming step in most cases. Therefore, one of the vCPUs was consistently underutilized during the trimming phase (Figure 6). However, *kallisto* maximized use of both vCPUs during most of its processing; it also consistently used all available RAM. Disk usage was heaviest while the FASTQ files were being unpacked, at the end of quality trimming, and at the beginning of *kallisto* execution. Because the data files were transferred to the VMs before our software container began executing, we detected no network activity (as expected).

**Figure 6.**
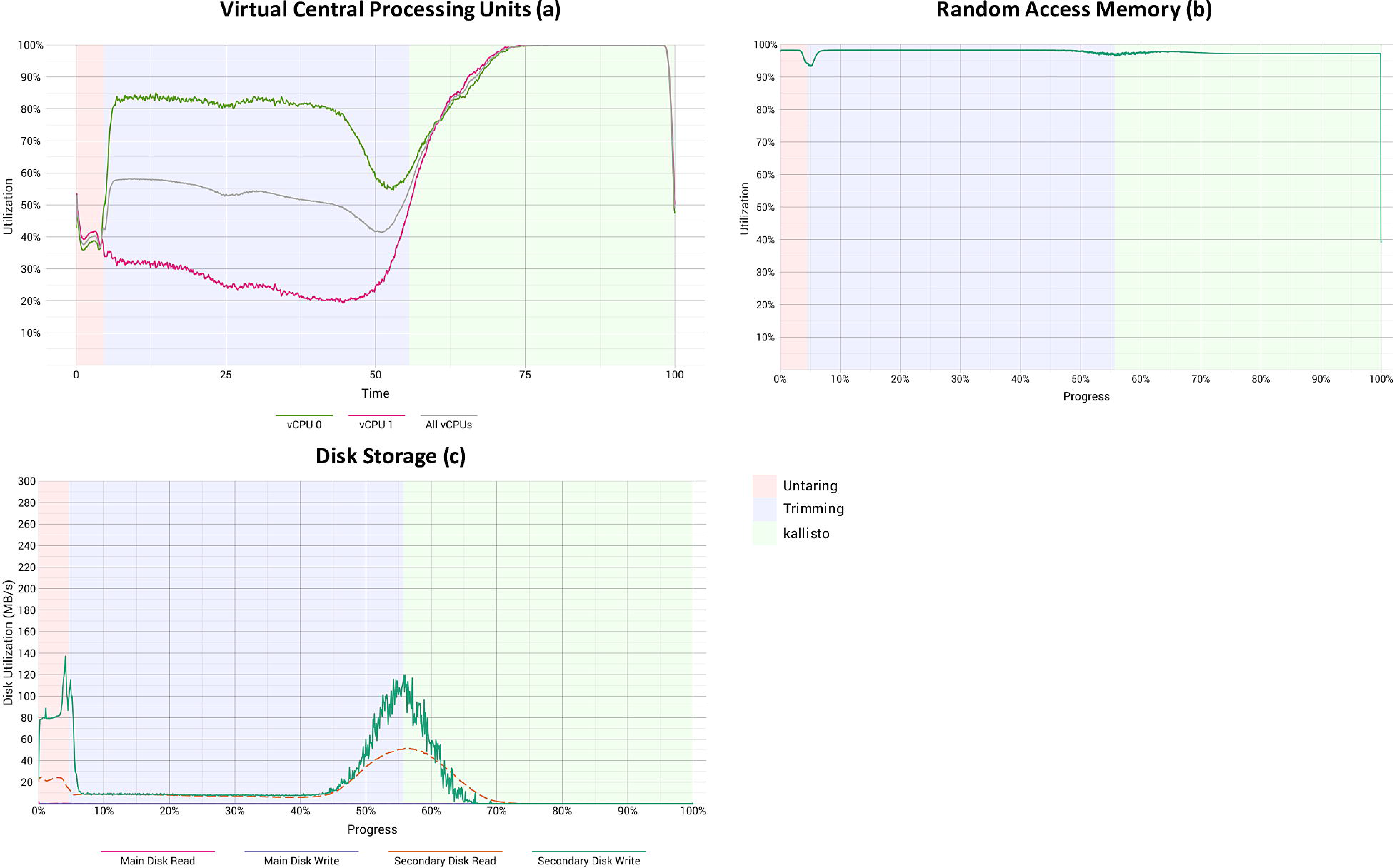
Computational resource utilization while TCGA breast and lung samples were processed using a preemptible-node configuration. These graphs show the (a) percentage of user and system vCPU utilization, (b) percentage of memory usage, and (c) disk activity. The “main” disks had 10 gigabytes of storage space and stored operating-system files. The “secondary” disks, which stored all data files, had 350 gigabytes of space. The background colors represent the computational tasks shown in Figure 4. We were unable to collect performance metrics for preliminary tasks, such as file localization, because these tasks were not performed within the software container. Each graph summarizes data observed across all 811 TCGA lung and breast samples. Because there was typically only one pair of FASTQ files per sample, quality trimming could not be parallelized; therefore, we used only 2 vCPUs per sample.

Lastly, we applied our software container to the remaining 9,562 TCGA samples, which ranged from 0.27 to 27.69 gigabytes in size. In total, raw sequencing data for the TCGA samples were 64.47 terabytes in size and consumed a total of 30,795.2 vCPU hours. Overall, the cost to process these samples was $1,065.49 ($0.09 per sample; see Tables 1-2). These costs are far lower than any prior cloud-based project of which we are aware.

We have deposited the processed data—as read counts and transcripts per million—in tabular format and in Google’s BigQuery database (see https://osf.io/gqrz9).

**Table 2:** Summary of cost information.

| Expense category<sup>a</sup> | CCLE<br>(cluster) | CCLE<br>(preemptible) | TCGA<br>(breast, lung) | TCGA (other<br>cancer types) |
| --- | --- | --- | --- | --- |
| Data transfer | \$0.02 | \$0.08 | \$0.10 | \$0.70 |
| Data storage | \$111.27 | \$25.48 | \$66.10 | 373.08 |
| Computing nodes | \$605.16 | \$338.09 | \$95.75 | \$520.80 |
| Miscellaneous | \$0.07 | \$0.57 | \$1.47 | \$8.12 |
| *Total cost* | *\$716.52* | *\$364.22* | *\$163.42* | *\$902.07* |
| *Cost per sample* | *\$0.767* | *\$0.390* | *\$0.090* | *\$0.094* |

<sup>a</sup> We obtained cost information for the Google Cloud Platform and assigned expenses to high-level categories. All values are in US Dollars (rounded to the nearest cent). All TCGA samples were processed using preemptible VMs. There is no charge for the time required to spinup VMs, pull images, localize files, or delocalize files.

## Discussion

The size of biomolecular data will continue to increase at a dramatic pace over the coming years^31^. To devise solutions for managing and processing these data, the research community will likely benefit from working with industry partners^43^, including cloud-computing providers^44–47^. Here we describe lessons learned as we quantified transcription levels for 12,307 samples from CCLE and TCGA. We hope these insights will be useful both to the academic community and to industry partners.

Our analyses were not exhaustive. Indeed, further optimization may yield better performance and lower cost than we observed. However, we have demonstrated that raw RNA-Sequencing data can be preprocessed, aligned, and quantified for pennies per sample. Furthermore, to enable others to reproduce our findings, we have provided all source code that we used to process the samples, to collect performance metrics, and to create the figures in this paper (see Methods). Our workflow can be used to process additional RNA-Sequencing samples available via the NCI Cloud Pilots or to process samples that have been sequenced by individual labs. With minor modifications to our pipeline, different reference genomes or algorithms could be applied to these data. We hope these resources will serve as examples to other researchers who wish to start using cloud environments.

Initially, we processed the CCLE samples using a cluster-based container engine, which is loosely comparable to cluster-computing environments commonly used at academic institutions. A disadvantage of this configuration is that the entire container engine needs to remain available while any sample remains to be processed. It may be possible to mitigate this limitation by processing the larger samples first (as we did); however, if any error occurs and sample(s) must be re-added to the queue, cost efficiency will decrease. When we used preemptible VMs to process the same samples, we did not need to pay for excess computing power and paid 50-70% less per computing node. Thus even though some samples were preempted and had to be reprocessed, the overall cost was much lower. We recommend this option as a way to reduce expenses.

Due to the way our workflow is structured, it was necessary to completely reprocess samples that had been preempted. An alternative solution would be to use persistent disk storage to store intermediate data files that result from individual processing steps within the workflow. When a sample is preempted, the workflow could check for these intermediate files and resume processing at the latest checkpoint. This approach might reduce overall processing times, but it would require users to rent additional storage space and thus might offset savings gained from reduced processing times. Yet another solution would be to split the workflow into smaller segments and execute the workflow in stages; again, this would require additional persistent storage, and the VMs would need to be deployed repeatedly, a step that can take several minutes at a time. Although such approaches may provide advantages in some cases, we favor simplicity, especially in situations where preemption occurs infrequently.

Our performance-profiling data demonstrate that computational resources are often underutilized by bioinformatics tools. For example, in our pipeline, vCPUs were used unevenly and often not at full capacity. Although it is unreasonable to expect that all computational resources can be used at full capacity at all stages of data processing, opportunities exist for tool developers to examine ways to optimize the use of these resources.

Due to the rapidly changing landscape of next-generation sequencing analysis, it is extremely beneficial for cloud providers to provide access to raw data. Until recently, raw sequencing data from CCLE and TCGA were available via the CGHub service^48^. Users who desired to compute on these data were required to bring “data to the tools,” whereas in cloud-computing environments, users can take their “tools to the data”^49^. In the case of CCLE, raw data were available in BAM format, which required us to convert the files to FASTQ format. In addition, we trimmed the reads for quality control. Together these steps took 70-75% of the processing time per sample (Figures 1 & S3). The TCGA raw data were available in FASTQ format; however, we needed to decompress and unpack *tar* files before trimming the reads (52% of the processing time). Accordingly, we recommend that alignment and quantification tools support both FASTQ and BAM input files—ideally, they would also support quality trimming. We also recommend that cloud providers store two versions of raw sequencing data: 1) compressed (but not packed) files that have been manipulated by no bioinformatics tool, and 2) files that have been sorted, trimmed, and compressed using standard tools. The former could be used by those who prefer to preprocess the data using alternative tools, whereas the latter could be used by all other users. Although storing these two versions of the raw data would increase disk-storage costs, it would likely decrease overall costs.

Cloud environments enable scientists to flexibly determine which resources are employed for a given research task. In contrast, academic computing environments are relatively fixed resources that may be oversubscribed, forcing individual researchers to wait long periods of time to process their data. At other times, these environments may be underutilized, thus reducing cost efficiency. In addition, because individual researchers typically do not have administrative privileges in academic computing environments, they may be limited in how they can customize their workflows. A central feature of our approach is the use of software containers, which enabled us to package, deploy, and monitor our software more readily. We anticipate that as containerization technologies continue to mature, they will be an integral part of cloud-computing workflows (and perhaps academic environments).

Commercial cloud-computing environments have been available for more than a decade. Sequencing and bioinformatics service providers commonly use cloud environments to apply bioinformatics tools to genomic data^50^. Recently, publicly funded, data-generating consortia, including projects funded by the National Cancer Institute, have begun to explore the benefits and challenges of providing centralized access to large compendia of genomic data. Our experiences confirm that the cloud has potential to make it easier to apply custom workflows to sequencing data at a modest price. Accordingly, we believe cloud computing will play an increasingly important role in cancer research. We encourage funding agencies and research institutions to consider implications of cloud computing on research budgets. Whereas academic-computing infrastructure has traditionally been funded by indirect costs, individual researchers may be required to categorize cloud computing as direct costs. Efforts such as The NIH Commons have begun to address this issue^51^. Finally, we encourage university training programs to place an emphasis on teaching students how to use cloud-computing resources.

## Methods

### Cluster-based configuration

The Google Cluster Engine allowed us to control hundreds of dedicated computing nodes with one central node. This central node used *Kubernetes^37^* to manage resources and a queue. Using a YAML^52^ configuration file, we submitted each RNA-Sequencing sample to *Kubernetes*, which then scheduled the samples to execute on the worker nodes. For each sample, *Kubernetes* deployed the relevant software container to the node, copied data files to the node, executed the software container, and copied output files to persistent storage. In cases where execution failed, *Kubernetes* automatically restarted sample processing. By default, the Google Cluster Engine allocated a 100 GB disk to each computing node; however, due to the size of the raw data and disk-space requirements for sorting and converting to FASTQ, we mounted a secondary disk drive (300 GB) to each node. While processing the samples, two nodes experienced errors while mounting disk drives. We used a two-node cluster to finish processing the remaining samples.

### Preemptible-node configuration

We used the Google Genomics Pipeline service^53^ to control execution of tasks on preemptible computing nodes. As a preemptible node became available, the service 1) created a VM on the node, 2) deployed the relevant software container to the VM, 3) copied data files to the secondary disk drive, 4) executed the software container, 5) copied output files to persistent storage, and 6) destroyed the VM after the sample finished processing (or was preempted). To help facilitate this process, we used a software framework provided by the Institute for Systems Biology^54^. Via a command-line interface, this framework facilitated the process of submitting samples to the Google Genomics Pipeline for processing. In addition, the framework monitored each sample’s status and resubmitted samples that had been preempted (at 60-second intervals).

### Software containers

We created three different Docker containers to house the software required to process each combination of data source and cloud configuration. The first container was used by the Google Cluster Engine to process BAM files from the CCLE samples. The cluster-based configuration required us to use the container to copy input files to and from the computing nodes. After copying the files, the container used *Sambamba* (version 0.6.0)^55^ to sort the BAM files by name and then *Picard Tools* (version 2.1.1, SamToFastq module)^56^ to convert the BAM files to FASTQ format. In accordance with *kallisto*’s documentation, we used the “OUTPUT_PER_RG” flag in *Picard Tools* to ensure that paired-end reads were placed in separate output files. The FASTQ files were then used as input to *kallisto* (version 0.43.0)^35^, which pseudoaligned the reads to the GENCODE reference transcriptome (version 24)^57^ and quantified transcript-expression levels. Based on the *kallisto* authors' recommendation, we used 30 bootstrap samples; we also used the “--bias” flag to account for variance and bias. We used the parallelization features in *Sambamba* and *kallisto* to enable faster processing.

The second software container is similar to the first but was modified for use with the Google Genomics Pipeline service. Because this service handles copying the data files between Google Cloud Storage and the computing nodes, these tasks were not performed by the container. We also added a read-trimming step using *Trim Galore!* (version 0.4.1)^58^, a wrapper around *Cutadapt* (version 1.10)^59^. This tool trims adapter sequences and low quality bases/reads. To process multiple FASTQ files (or pairs of FASTQ files for paired-end reads) in parallel, we used *GNU Parallel* (version 20141022)^60^.

The third software container was designed specifically for the TCGA data. It extracts FASTQ files from a tar archive (whether compressed or not), performs quality trimming, and executes *kallisto*. Where applicable, it uses the *pigz* tool (version 2.3.1)^61^ to decompress the input files in parallel.

All three containers use the *sar* module of the *sysstat* program (version 11.2.0)^62^ to log each machine’s vCPU, memory, disk, and network activity throughout the course of data processing. The containers copied these data to persistent storage, prior to the job’s completion. We changed the time of each entry in the logs to a corresponding percentage of total job time, to allow the activity metrics to be summarized consistently across all jobs.

### Code availability

We compiled an open-access repository (https://osf.io/gqrz9) that contains all the scripts we used to construct the Docker containers and to process samples on the Google Cloud Platform. The repository also includes summarized performance data and scripts that we used to generate the figures in this manuscript.

## Acknowledgements

SRP thanks Brigham Young University for startup research funds. PJT thanks the Simmons Center for Cancer Research at Brigham Young University for a summer fellowship that enabled him to perform this study. We thank the National Cancer Institute for providing cloud credits via the ISB Cancer Genomics Cloud pilot. We thank Sheila Reynolds and Abigail Hahn from the ISB for providing technical support and feedback on our manuscript. We thank researchers at the Broad Institute and The Cancer Genome Atlas Consortium who released data to the public. Most importantly, we thank the patients who have donated tissue samples and consented to data sharing.

### Author Contributions

PJT developed the software pipeline, processed all data, designed analyses, created all figures, and wrote the paper. SRP developed the project idea, designed analyses, and wrote the paper.

